# Length control of filamentous structures in cells by the limiting pool mechanism

**DOI:** 10.1101/075655

**Authors:** Lishibanya Mohapatra, Thibaut J. Lagny, David Harbage, Predrag R. Jelenkovic, Jane Kondev

## Abstract

How the size of organelles in cells is controlled despite a constant turnover of their constituent parts is a central problem in cell biology. A general mechanism has been proposed based on the idea that an organelle grows by self-assembly of molecular subunits that freely diffuse in the cytoplasm. Assembly continues until the available pool of subunits is depleted to the point when the stochastic addition and removal of subunits is balanced, leading to a structure of well-defined size. Here we focus on length control of multiple filamentous structures in cells, such as actin cables and flagella. Using queueing theory and computation we show that the limiting pool mechanism leads to three different phases of assembly, starting with a rapid growth phase when all filaments quickly accumulate a large number of available subunits. Then, the slower growing filamentous structures enter a disassembly phase as they gradually lose all of their subunits to the faster growing structures. Finally, when multiple, equivalent fast-growing filaments are present, their lengths undergo protracted diffusive dynamics due to the stochastic swapping of subunits between them. This eventually leads to a broad, power-law distribution of filament lengths in steady state. Our findings demonstrate that the limiting-pool mechanism is incapable of controlling lengths of multiple filamentous structures that are assembled from a common pool of subunits, and at best, can produce only one filament of a well-defined size. Overall, our theoretical results reveal physical limitations of the limiting-pool mechanism of organelle size control.

**Significance Statement:** What determines the size of organelles in cells is a classic problem in cell biology. Recent experiments on mitotic spindles, and nucleolus have singled out the limiting-pool mechanism of size control. As these structures assemble, they deplete a finite pool of subunits present in the cell, thereby reducing the rate of subunit addition. Eventually the stochastic addition and removal of subunits are balanced and a well-defined size is achieved. We find that, while the limiting-pool mechanism does control the size of an individual structure, it fails when multiple structures are competing for the same pool of subunits. In that case we predict large size fluctuations and that the fastest growing structure takes up practically all the subunits from the pool.

## Introduction

Cells consist of organelles whose size is often matched to the size of the cell. The scaling of the spindle size with the size of the cell in a developing embryo is a classic example (1). How these organelles are assembled and maintained to have a specific size is a question that is still not well understood. A simple idea that seems to provide the answer in many but certainly not all cases, is that the cell maintains a limiting pool of a molecular component that is required for assembling the organelle. In such a case, size control is simply achieved by the organelle growing until the limiting pool is depleted to the point when the rates of assembly and disassembly are matched. In this paper we theoretically investigate the limiting pool mechanism for controlling the length of filamentous structures in cells. In particular, we identify its limitations in controlling size when two or more structures are assembled from the same building blocks.

The cell cytoskeleton contains a number of micron-scale filamentous structures which are maintained at a constant length suggesting that their form is related to their function. Examples include mitotic spindles, flagella, actin patches and actin cables(2–5). Large changes in their size often result in significant deviations from their normal physiological functions. For example, In yeast cells intracellular transport is disrupted if actin cables overgrow and buckle(5). In fact, experiments have shown that when filamentous structures are cut to a smaller size, they often grow back to their physiological length suggesting that the length is under tight control(4).

Most cellular filamentous structures are composed of actin filaments and microtubules, which in turn are composed of actin monomers and tubulin dimers, respectively. These subunits undergo constant turnover as they are stochastically added and removed from the structure. The key question then is how the balance between addition and removal of molecular building blocks is achieved so as to generate a structure of a well-defined length. Another related question is how different-sized structures, made of the same building blocks can coexist in the same cell. For example, actin cables and actin patches in yeast are made up of the same actin monomers, have different size, form, and function, yet co-exist in the same cytoplasm and exchange actin monomers(6).

The idea that a limiting pool of diffusible subunits plays a major role in the control of size has received considerable attention as it offers a simple mechanism of size regulation. A recent review summarized the experimental evidence for this mechanism controlling the size of diverse structures such as centrosomes, flagella, and the nucleus(1). More recently, *in vitro* studies used a microfluidic system to encapsulate cytoplasm from *Xenopus* egg extracts in small droplets and showed that spindle size is proportional to the droplet volume, thus suggesting that the amount of cytoplasmic material controlled its size(7). Another recent study showed inverse scaling of the size of nucleoli with cell size in a developing *C. elegans* embryo in which the number of subunits in the nucleoplasm was fixed, also consistent with the limiting pool mechanism(8). The idea of a limiting-pool is particularly appealing as it offers a simple size-control mechanism without the need for any additional molecular mechanism by which cells “measure” individual structure sizes and adjust assembly or disassembly rates accordingly.

The question that we address in this paper is whether the limiting pool mechanism is capable of maintaining multiple structures with a well-defined size, when the structures share a common molecular component that is limiting. We consider this general question in the context of a simple model of filament growth, from which we derive general conclusions about the limiting pool mechanism.

## Results

We consider the limiting-pool mechanism of size control in the context of a simple model where filaments grow from a fixed number of nucleating centers within the cell by stochastic addition of diffusing monomers. Monomers, whose number in the cell is fixed, also stochastically dissociate from the filament. The number of filaments is fixed by the number of nucleating centers, which can be a single protein or a protein complex, and which aid in the formation of the filament. An example is provided by formins which help assemble filamentous actin structures. Formins bind to the barbed end of an actin filament and capture (profilin bound) actin monomers from solution, which are then incorporated into the growing filament (see Figure 1A). Note that the model we consider is a significant departure from textbook examples of stochastic filament growth where every monomer in solution can serve as the site of new filament growth. In our case filament growth occurs only from nucleating centers.

We consider three different scenarios, one when there is a single nucleating center in a cell which contains a fixed number of monomers, the case of two identical nucleating centers, and of two distinct nucleating centers, which differ in the rates at which they incorporate monomers. An example of inequivalent nucleators is provided by the two different formins Bni1 and Bnr1 in budding yeast, which assemble actin filaments at different rates(9).

### A limiting-pool of monomers leads to a single filament of a well-defined length

First we consider the case of a single nucleating center, where a single filament is assembled by the addition and dissociation of monomers. The total number of monomers in the cell, *N*, is fixed, and each monomer can associate to a filament with growth rate *k*_*_, which is proportional to the number of free monomers in the compartment. Hence, for a single filament, its growth rate starts off as 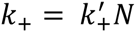 but as the filament grows, it decreases to 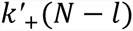 where *l* is the length of the filament in units of monomers. Note that the rate constant 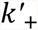 is obtained by taking the second order rate constant for monomer addition, which has units *M*^−1^*s*^−1^, and multiplying it by the volume of the cytoplasm within which the free monomers diffuse. The rate of growth is thus length dependent, and assuming a constant monomer dissociation rate, *k*_, it leads to a peaked distribution of filament lengths (Figure 1B).

**Figure 1:**
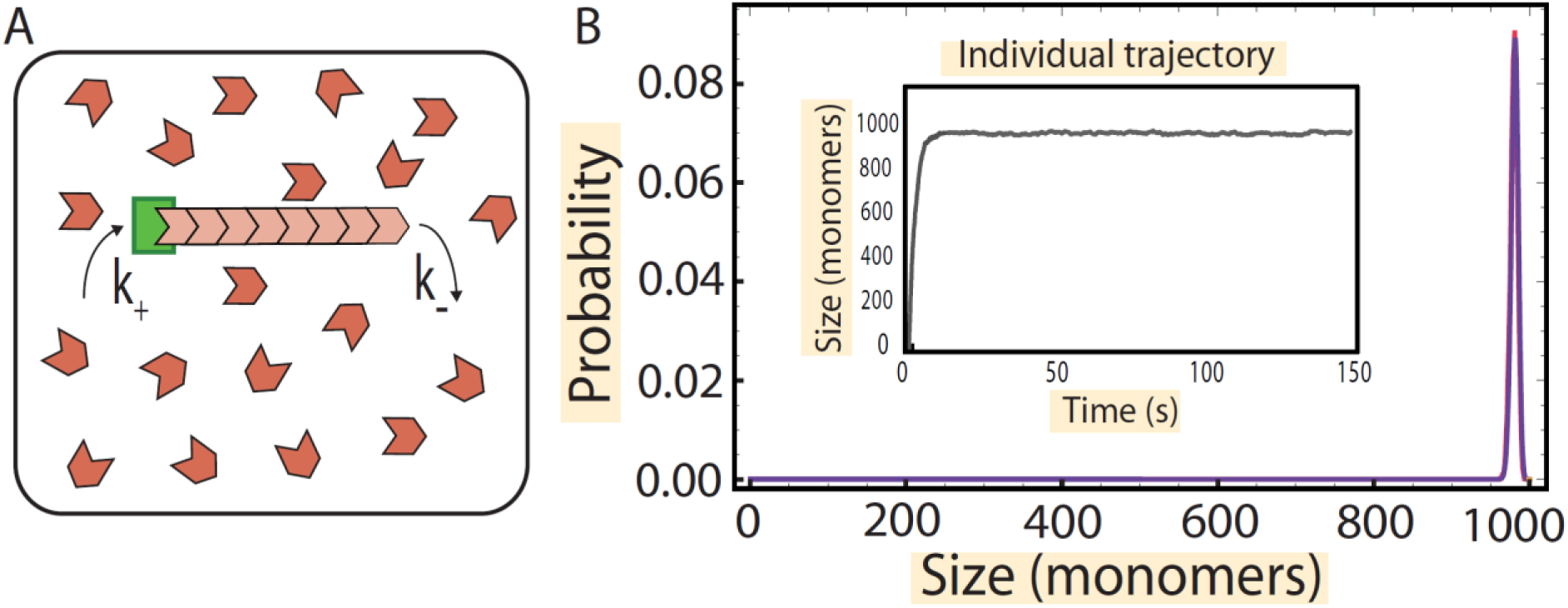
Growth of a filament from a single nucleating center in a limiting pool of monomers. (A) Schematic showing the growth of a single filament (pink) from a single nucleating center (green) in a pool of monomers (in red). (B) (Inset) Numerical simulation of the growth trajectory of a single filament from the nucleating center (gray). After a fast growth phase, the size attains a steady state. By considering several such trajectories, we compute the probability distribution of filament lengths numerically, and a peaked distribution of size is obtained (in red). The parameters used for simulations are 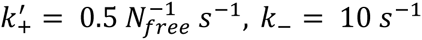 and *N* = 1000, where *N*_*free*_ is the number of free monomers in solution. The simulation results are compared with the analytical results (blue) obtained in SI, section 4.

In order to describe the dynamics of an individual filament, we model the growth and decay of the filament using the master equation formalism. The key quantity to compute is the probability, *p*(*l*, *t*), that the filament has a length *l* (measured here in units of monomers) at time *t*. The master equation describes the evolution of *p*(*l*, *t*) in time, by taking into account all the possible changes of the state (length) of filament that can occur in a small time interval Δ*t* (Figure 1A). The master equation for a single filament is (for *l* > 0)

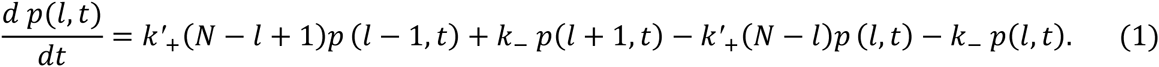

We compute the steady-state distribution of filament lengths by setting the left-hand side of the equation to zero and omit the time variable in *p*(*l*) to indicate the steady state nature of the distribution. We use detailed balance equations 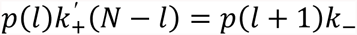 to obtain 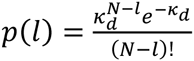 (SI, section 4), where 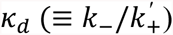 is a dimensionless dissociation constant for the chemical reaction of a monomer binding to filament which is obtained by multiplying the dissociation constant with the cell volume. For example, for actin cables in yeast cells we estimate 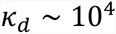 and *N* ~ 2 × 10^5^, using the measured concentration of actin in yeast (~ 10*μ*M) (10), the typical volume of a yeast cell (~ 40*μ*m^3^) (11) and the measured rates of association (11.6 *μ*M^−1^*s*^−1^) and dissociation (1.4 *s*^−1^) for binding of actin monomers to actin filaments (12) (See SI section 3.5 for the calculations). The mean and standard deviation of the distribution are given by 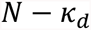 and 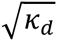 respectively. This distribution is very peaked, and the same is true for the distribution of free monomers, which in fact is very close to Poisson (See SI, section 3). Interestingly, the typical length of the filament is essentially given by the number of available monomers unless *k*’_=_ and *k*_ are fine-tuned to be close in value (See SI, section 3.1).

We used stochastic simulations to analyze the time evolution of the length distribution. We start with a filament of zero length growing from a single nucleating center and then follow the growth trajectory of the length of the filament in time as monomers attach and fall off. After some time, we observe the filament reaching a steady state (see Figure 1B inset), when the length distribution of the filaments no longer changes with time. The distribution extracted from these simulations matches the analytic results. The time scale over which the steady state is reached is of order 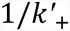 which can be understood as the time it takes *N* monomers to be taken up from the pool at a rate 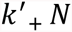 (see SI, section 2.1 for a more precise calculation).

### Two identical nucleating centers in a limiting pool of monomers produce filaments with large, anti-correlated fluctuations in length

Next we turn to the case of assembly of multiple filamentous structures competing for a common, limited pool of monomers. We begin by considering the simplest case of two nucleating centers that are growing one filament each (Figure 2A). The results obtained in this simple case are also found in the more general many-filament case, which is described in detail in the SI.

**Figure 2:**
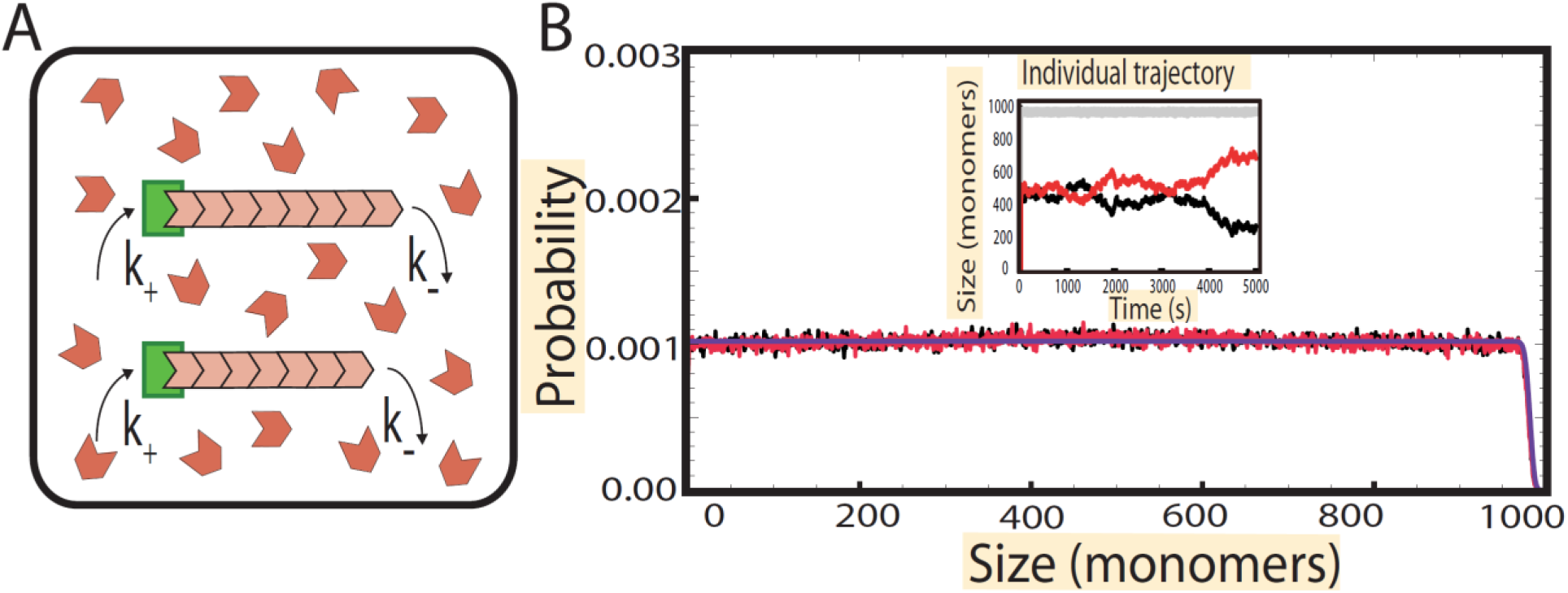
Growth of filaments from two identical nucleating centers in a limiting pool of monomers. (A) Schematic showing the growth of filaments (pink) from two nucleating centers (green) in a pool of monomers (in red). **(B)** (Inset) Numerical simulation of the growth trajectory of the two individual filaments (red and black) and the sum of their lengths (gray). After an initial growth phase where the two filaments grow roughly in unison and the total length of the two filaments reaches steady state, we observe anti-correlated fluctuations in the individual lengths. These fluctuations lead to individual filaments having a uniform distribution of lengths (red and black) in simulations; we compare these to results obtained analytically (blue) in SI, section 4.3. The parameters used for simulations are 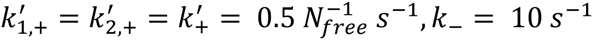 and *N* = 1000, where *N*_*free*_ is the number of free monomers.

In the case of two identical nucleating centers, each of the two filaments adds a monomer to it at the same rate 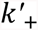. Therefore the growth rate of filaments starts off as 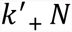, however as the two filaments grow, it decreases to 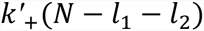, where *l*_1_ *l*_2_ and are the lengths of the two filaments in units of monomers. The rate of dissociation (*k*_) of monomers is identical for both filaments.

Since there are now two filaments in the cell, we characterize the state of the system with a joint probability distribution *p*(*l*_1_,*l*_2_, *t*), which satisfies the master equation

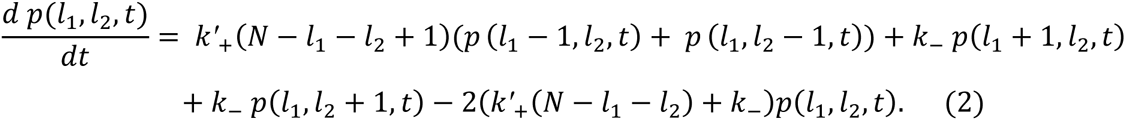

At steady state, *d p*(*l*_1_,*l*_2_,*t*)/*dt* = 0 and the joint probability distribution takes on the product 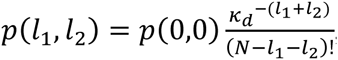 which is a result from queueing theory(13) (see SI, section 4). Here 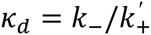 is once again the dimensionless dissociation constant. The distribution of lengths for a single filament, i.e., *p*_1_(*l*_1_) and *p*_2_(*l*_2_), can be obtained from the joint distribution by summing over all possible lengths of the other filament (*l*_2_ and *l*_1_, respectively); we perform these calculations explicitly in sections 4.1 and 4.3 of the SI.

An approximate formula for the distribution of lengths for a single filament in steady state can be derived from a simple argument, which also sheds light on the physics of the problem. In steady state the average number of free monomers left in solution is 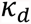 and the distribution of free monomers is very close to Poisson with standard deviation 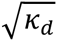, implying that it is very narrowly concentrated around its mean 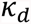. (The Poisson approximation to the free monomer pool was previously derived (14) by assuming that the probability of a filament having zero length is zero; we rigorously justify this approximation in SI, section 4.1) Hence, the total number of assembled monomers is to an excellent approximation a constant 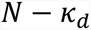. Therefore, by symmetry, every possible configuration of filament lengths that satisfies 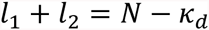 is equally likely. The number of such confgurations is 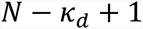 and therefore

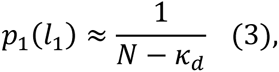

i.e., it is uniform on the interval 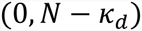. This approximation is very accurate except in a very narrow interval of lengths of order 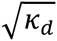 when *l*_1_ is close to 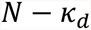, where it decays rapidly to zero. This effect is due to the Gaussian fluctuations of the free monomer pool around its mean. This simple argument can be further extended to an arbitrary number of filaments and is validated by our exact calculations (see SI, sections 1.1 and 4.1).

We also used stochastic simulations to investigate individual growth trajectories for the two filaments and compared the results to our analytical steady-state distributions *p*_1_(*l*_1_) and *p*_2_(*l*_2_) (Figure 2B). Once again, we follow the stochastic trajectory in time of the length of each individual filament. Initially, both filaments grow in unison (subject to small fluctuations) until their combined length reaches the steady state total length 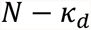. After this rapid growth period the individual trajectories of the filaments diverge, as one grows the other shrinks (Figure 2B inset). These anti-correlated fluctuations in length persist indefinitely and eventually become of order 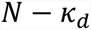. At the same time the length distributions for each filament,*p*_1_(*l*_1_) and *p*_2_(*l*_2_), settle into their steady state values, Equation 3.

These dynamics are illustrated in Figure 3A, where we plot *p*_1_(*l*_1_, *t*) obtained from stochastic simulations. At first the mean filament length increases until the free monomer pool (or equivalently the total filament length) reaches its steady state (black curves). The time scale for this is of order 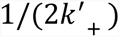, same as in the case of a single filament. After that, we observe the widening of the distribution at constant mean (blue curve) which eventually becomes uniform (red curve) at the longest times, of order *N*^2^/*k*_ (SI, section 2.3). Thus, we reach the somewhat surprising conclusion that the limiting-pool mechanism does not control length in the case of two structures competing for the same monomers. In fact, this remains true when more than two identical structures are present, as explained in the Discussion section.

**Figure 3.**
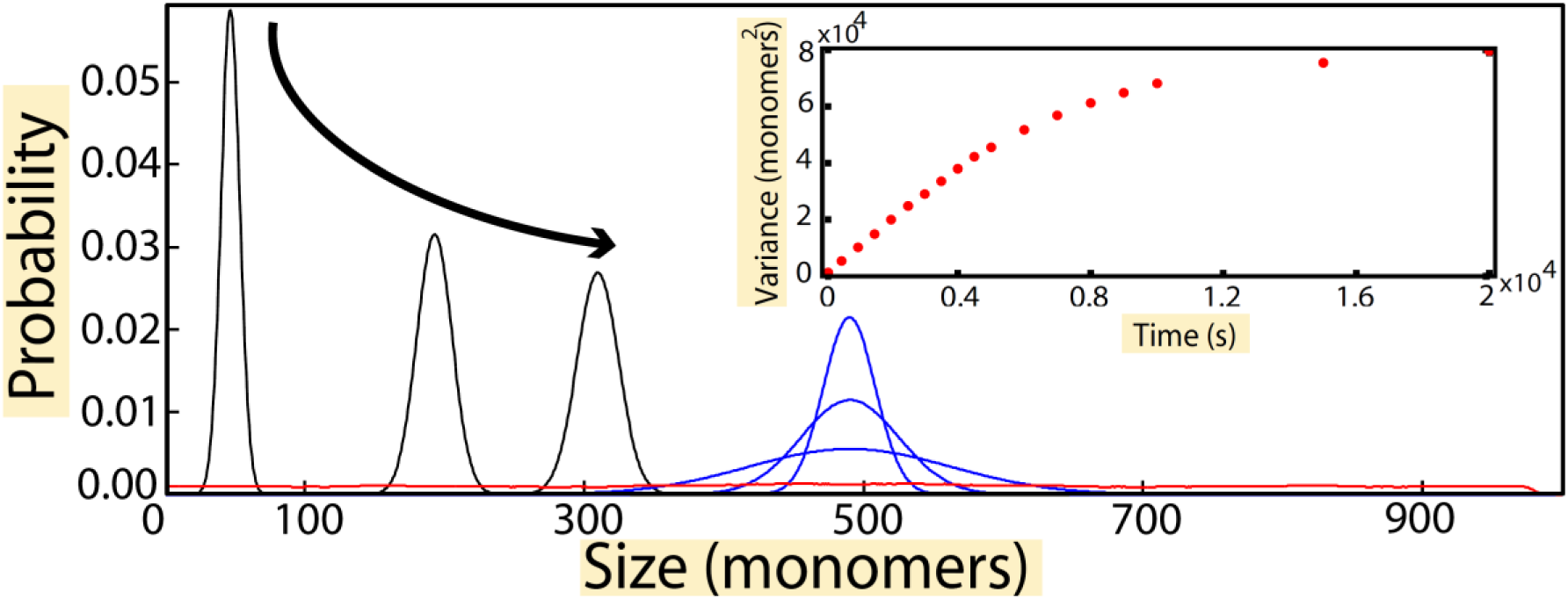
Evolution of probability for individual filament lengths in the case of identical nucleating centers. In the initial growth phase (black), the distribution of individual filament lengths starts off as a peaked distribution whose mean increases, until the total length reaches a steady state. The growth phase is followed by the slow phase of monomers swapping which leads to anti-correlated fluctuations are seen in the individual trajectories, which is translated into the increase in the width of the filament length distributions while the mean stays unchanged (blue). The final, steady state distribution (red) is flat. (Inset) Plot of variance of the distribution (*σ*^2^) vs. time of simulation. The variance initially increases linearly with time, but later saturates. Time was varied from 0 — 20000 *s* in steps of 100 *s*. From the slope of the linear part plot of, we find a diffusion constant *D* = 5*monomer*^2^/*s* using the relation *σ*^2^ = 2*Dt*. The parameters used for simulations were 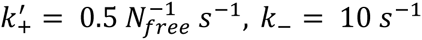, *N* = 1000 where *N*_*free*_ is the number of free monomers.

Once the total length of the two filaments has reached a steady state, any monomer that come off one filament is rapidly taken up by the other filament. This is a consequence of the fact that in steady state the total number of monomers in the two filaments is narrowly distributed around its mean value. We therefore expect individual filament lengths to exhibit diffusive dynamics. We demonstrate this explicitly in the inset to Figure 3B where the variance, *σ*^2^, of the length distribution of an individual filament is seen to increase linearly with time until it eventually saturates. Given the diffusional law *σ*^2^ = 2 *Dt* the initial slope of the *σ*^2^ curve in Figure 3B gives the diffusion constant *D*. The diffusion constant can be computed by taking into account that when the total length reaches steady state monomers exchange between the two filaments with a rate set by *k*_. A detailed analysis of all possible exchange processes while accounting for their probabilities leads to the exact result *D* = *k*_/2 (see SI, section 2.3), which we also checked with stochastic simulations.

### Two inequivalent filaments assembling in a limiting pool of monomers results in only one filament having a well-defined length

Next, we consider the case of two nucleating centers with different elongation rates for filaments in a common pool of monomers (see Figure 4A). As in the example of two different formins in yeast, monomers are stochastically added to the two filaments at different rates: 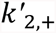 and 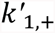. These rates start as 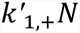 and 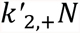 but as the two filaments grow they decrease to 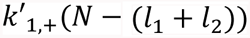 and 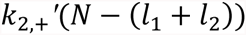 respectively, where *l*_1_ and *l*_2_ are the individual lengths of the two filaments in units of monomers. The rate of dissociation, *k_*, of monomers is assumed identical for both filaments.

**Figure 4:**
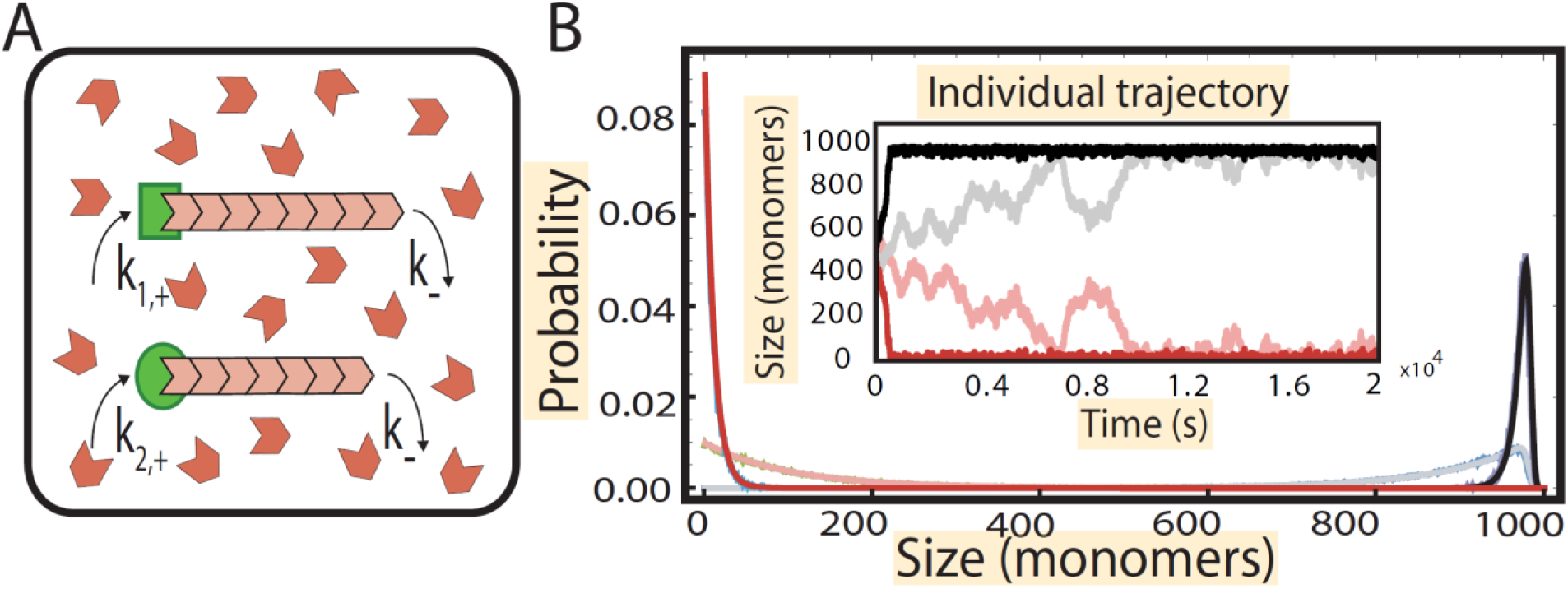
Growth of filaments from two inequivalent nucleating centers in a limiting pool of monomers. (A) Schematic showing the growth of filaments (pink) from two distinct nucleating centers (green) in a pool of monomers (in red). (B) (Inset) Numerical simulation of the growth trajectory of the filaments from the nucleating centers. Shown are trajectories for 10% difference in elongation rate (dark) and 1% difference (light). After a growth phase, where both filaments accrue monomers, the faster growing filaments attains a steady state by taking up most of the free monomers, while the slower filament shrinks. The faster growing filament attains a peaked distribution of size (black) and the slower one attains a geometric distribution (red). The parameters used are 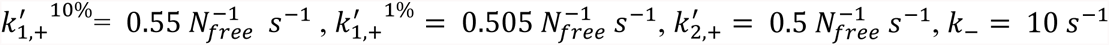 and *N* = 1000. The simulations are overlaid on the results obtained analytically (blue and green) in SI, section 4.3.

As before, the joint probability distribution *p*(*l*_1_, *l*_2_, *t*) satisfies a master equation, but now with different assembly rates for the two filaments, namely

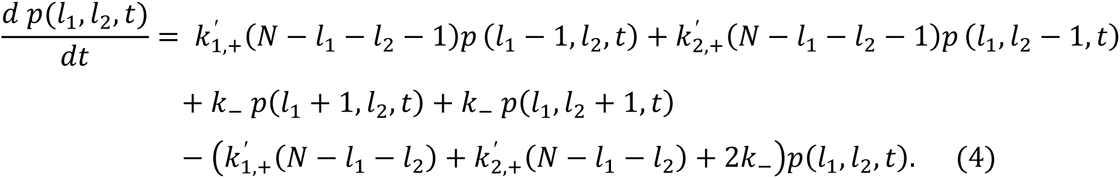

The steady state joint distribution of filament lengths is again given by a product form, from which the distribution of lengths of each individual filament can be computed. The exact calculation is in the SI and here we provide a simple intuitive argument.

For concreteness, let us assume that the growth rate of first filament is larger than the second, i.e., 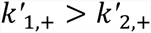 (or equivalently, 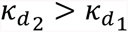, using 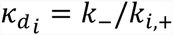 for the dimensionless dissociation constants). Then over a long period of time, a larger number of monomers will join the first, fast assembling filament. Since the detachments rates are the same, the first filament will accumulate most of the monomers. Hence, *p*_1_(0) ≈ 0, implying that average rate of monomers leaving the first filament is *k*_(1-*p*_1_(0)) ≈. Next, by equating this rate to the rate at which monomers attach to the first filament, i.e., 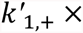 (average number of free monomers), we find that the average number of free monomers is 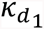, and, as in the case of equivalent filaments, we expect the steady-state distribution of the number of free monomers to be Poisson. From this result, we can derive the distribution of lengths for the second, slower growing filament from the detailed balance equations 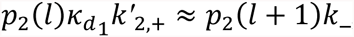, which leads to the conclusion that the lengths for the second filament are distributed geometrically, i.e.,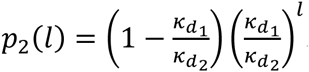. Then the first filament is virtually unaffected by the presence of the second and the distribution of its lengths is peaked, as in the case of a single filament. The approximate average length of the first and the second filament are given by 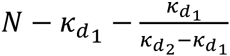 and 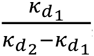, respectively (see SI, sections 1.2 and 4.2).

Once again, in order to develop intuition about the assembly dynamics we used stochastic simulations to follow the growth of the two filaments from the two nucleating centers each starting from zero length (Figure 4B, inset). Just like in the case of equal nucleating centers, we observe that there is an initial, fast growth scale which occurs over a time of the order 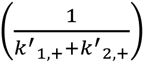. After the growth phase, we observe that filament with a higher growth rate grows to a steady state characterized by a large length, whereas the other filament shrinks to zero. When the two elongation rates are close in magnitude then we see for a while anti-correlated fluctuations of lengths similar to what we observed in the case of identical nucleating centers. These fluctuations happen over a time scale 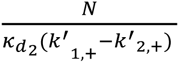,i.e., over a time that is of order *N*, and eventually subside as the systems settles into a steady state with practically all monomers taken up by the faster growing filament. This is illustrated in the plots of length distributions for the two filaments, where for the fast-assembling filament we observe a distribution sharply peaked around the mean while the slower assembling filament is characterized by a geometric distribution of filament lengths peaked at zero; see Figure 4B. We find wider distributions when the elongation rates are numerically close to each other, which results from the increased fluctuations of the filament lengths. Comparison of the analytical solution of Equation 4 for *p*_1_(*l*_1_) and *p*_2_(*l*_2_) (see SI, section 1.2 and 3.2) to the simulation data serves as a stringent test of the simulation procedure (Figure 4B).

To summarize, when considering two nucleating centers with different rates of filament assembly, the one with the higher rate wins and produces a filament of well-defined length, while the other filament does not stably form.

## Discussion

The limiting pool mechanism of size control has been implicated in assembly of a number of different cellular structures in different cell types (1). In order to understand the quantitative features of this general size-control mechanism we considered a simple model of stochastic growth of filamentous structures from a limiting pool of diffusing monomers, where the number of filaments is determined by the number of nucleating centers at which they assemble. While we found that the limiting-pool mechanism is able to precisely control the length of a single filament, it is unable to control individual filament lengths when two filaments are assembled from the same monomer pool.

Namely, when the two nucleating centers produce identical filaments, we found that the total length of the two generated filaments reaches a well-defined value in steady state, however their individual lengths are characterized by large fluctuations, which are only limited by the total number of monomers. In the presence of two non-identical nucleating centers which assemble filaments with different growth rates, the one with the higher growth rate “wins”, i.e., it takes up most of the available monomers.

### Assembly of many filaments

Thus far we have only considered two filamentous structures competing for the same pool of monomers. In cells though it is common to observe multiple structures made from the same pool of monomers. For example, the number of actin cables in budding yeast is about ten or so, while the number of actin patches is of order ten to a hundred.

The key results described above for two filamentous structures readily carry over to the case of multiple filaments. More specifically, for any finite number of filaments starting at zero length, growth of all filaments is strongly favored in the initial phase of assembly, and all the filaments quickly reach lengths of order *N* (the total number of monomers) over a time scale which is independent of *N*. Then, in case of inequivalent elongation rates, the slower growing filaments gradually lose monomers and diminish in length to a small, geometrically distributed length. The duration of this phase is of order *N*. Furthermore, at the end of this phase practically all the monomers are taken up by the fast growing filaments, and the number of free monomers approaches a Poisson distribution with a mean equal to the smallest (dimensionless) dissociation constant among all the filaments. This then implies that the total number of monomers assembled into the fast growing filaments will have a well-defined size characterized by a peaked distribution. The lengths of individual fast-growing filaments then undergo a protracted diffusive dynamics on time scales of the order *N*^2^. These dynamics are generated by the stochastic swapping of monomers between individual filaments, and eventually lead to a broad, power-law distribution of filament lengths in steady state. Interestingly, we find that if the number of fast growing filaments is of the order of *N*, then in the limit of large *N* the resulting length distribution is geometric. This theoretical result might provide a link between two recent experiments on mitotic spindles, one which showed that the spindle size is controlled by a limiting pool mechanism (7) and the other that found that individual microtubule lengths within the spindle are geometrically distributed (15).

To illustrate our general results regarding assembly of many filaments from a common pool of monomers we show in Figure 5 the different phases of assembly using an example of six filaments with three growing at the same, fast rate and three taking up monomers more slowly. (See SI for detailed calculations pertaining to the multi-filament case.)

**Figure 5:**
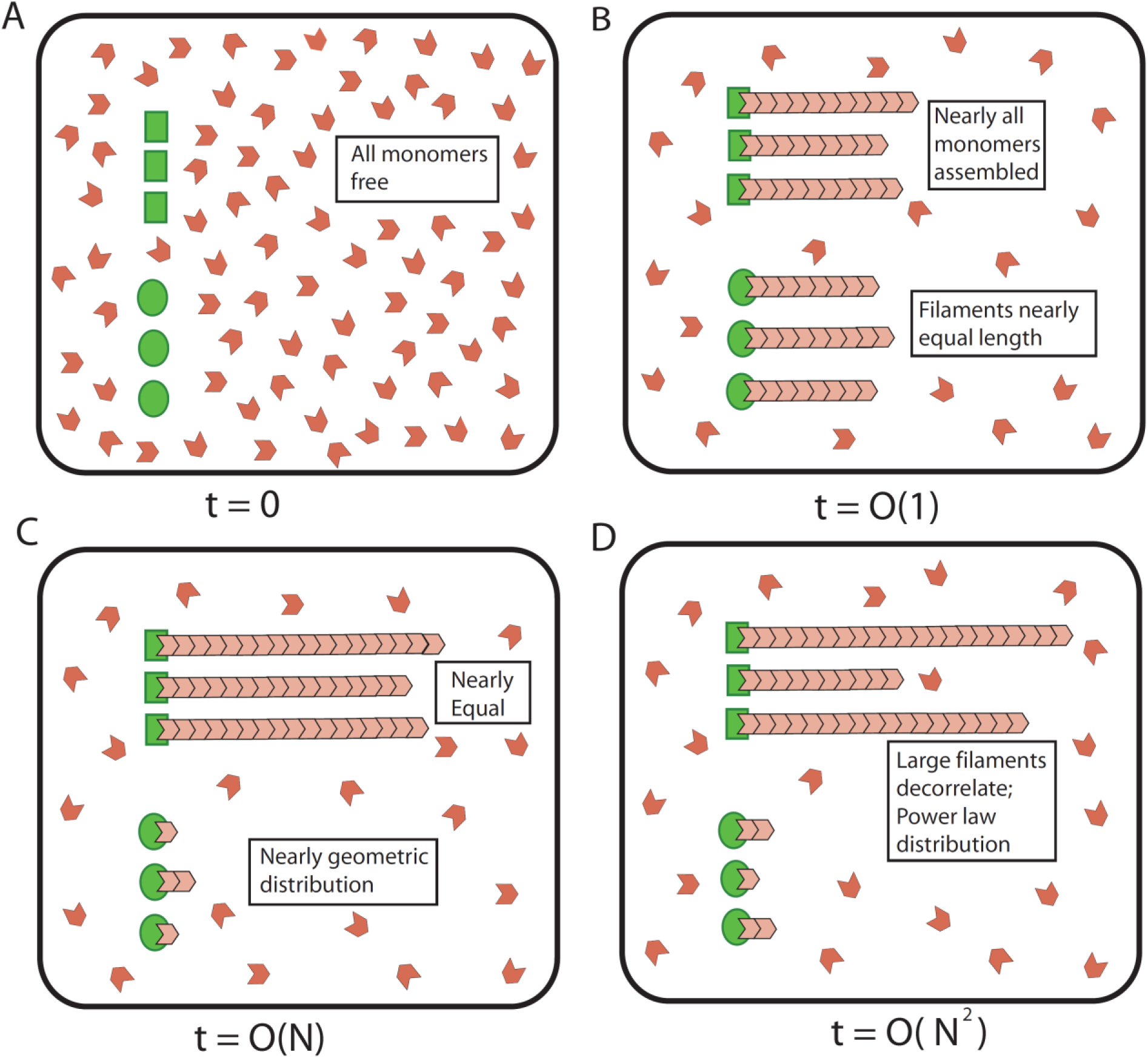
Growth dynamics of six filaments from two types of inequivalent nucleating centers. (A) Schematic showing six nucleating centers in a pool of monomers where the first three nucleate filaments that grow at a high association rate *k*′_*h*,+_, and the remaining three at a slower rate *k*′_s,+_(*k*′_*h*,+_ >*k*′_*s*,+_); all filaments disassemble with the same rate *k*_.(B) After a rapid assembly phase lasting approximately 1/(3(*k*′_*h*,+_/+*k*′_*s*,+_)), the fast and slow growing filaments reach the average length of *Nk*′_*h*,+_/(3(*k*′_*s*,+_)) and *Nk*′_*s*,+_/(3(*k*′_*h*,+_ + *k*′_*s*,+_)), respectively. (C) After this rapid assembly phase, the slower growing filaments slowly decrease to nearly zero length over a period of time of order 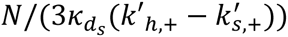, where 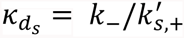 is the (dimensionless) dissociation constant for the slow growing filaments. At the end of this phase, the three slow growing filaments are geometrically distributed with parameter 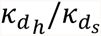 the free monomer pool approaches a Poisson distribution with mean 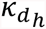 and nearly all the monomers,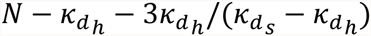, are taken by the fast growing filaments, and are equally distributed among them. (D) Finally, the sizes of the three largest filaments decorrelate on a slow, diffusion time scale which is of order *N*^2^, during which the monomers randomly exchange between these large filaments.

### Assembly of higher dimensional structures

From the preceding analysis we see that assembling multiple filamentous structures from a finite and limiting pool of monomers does not lead to precise size control of individual structures without additional control mechanisms. This stems from the inherent reversibility of the assembly process, which allows the monomers to exchange between the different structures, resulting in either very small sizes, or high variability described by a power-law distribution of sizes for individual structures. Since the key ingredient, namely the reversibility of stochastic assembly, is not limited to linear structures, similar phenomena are likely to occur in two or three-dimensional structures assembled from a common and limiting pool of subunits.

### Experimental tests of the limiting-pool mechanism

Our study makes a number of predictions that can be used to test the limiting-pool mechanism. In case of a single filamentous structure assembled from a pool of monomers, the steady state distribution of filament length (SI, Section 3) can be tested in experiments in which the total number of monomers is tuned. This can be achieved, for example, by using the microfluidic approach described in (7). Interestingly, we observe that the mean length (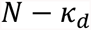) of the filament depends on the total number of monomers whereas the variance 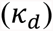 does not. This result can be used as a stringent test of the limiting-pool mechanism of size control.

Secondly, if there are multiple identical structures being made from a common pool of monomers, we predict the existence of anti-correlated fluctuations of individual filament sizes over time. For the two filament case, we predicted that these fluctuations will be observed at time scales of order *N*^2^/*k*_, which can also be tuned by controlling the total number of monomers. An experiment with two inequivalent filaments assembling from a common monomer pool should also reveal the time scale of order *N*, during which the slower assembling filament lose the monomers they quickly accumulated in the initial growth phase.

For example, in *C. elegans* cells, two nucleoli are assembled from the same pool of nucleoli particles and they grow equally in size up until cell division, which happens after about 20 minutes (8). However, if we were to engineer cells with longer cell cycles it might be possible to see the predicted anti-correlated fluctuations in individual nucleoli sizes, assuming the size of these structures is controlled by a limiting pool mechanism alone.

Another example where in vivo experiments can be used to test our predictions is provided by fission yeast cells. These cells have two different types of actin structures, namely cables and patches which are assembled by different nucleating factors (formins and the Arp2/3 complex, respectively) (16, 17). Recently it was shown that it is possible reduce the number of patches in yeast cells by over-expressing profilin, which is an actin-binding protein having two specific effects: it significantly favors the formation of cables by increasing the elongation rates of formin-nucleated filaments and it inhibits Arp 2/3 mediated branching and hence represses the formation of patches(16, 17). Thus by regulating the level of profilin either formin or Arp 2/3 generated structures will take up most of the available pool of actin monomers. This observation is consistent with our calculations since we find that when two structures are competing for the same subunit pool the one that assembles faster takes up practically all subunits. Still, further experiments need to be performed in which size distributions of different structures are measured to quantitatively test predictions of the limiting monomer pool model.

Finally, the limiting pool mechanism has also been invoked to explain size regulation of flagella in *Chlamydomonas,* which are microtubule filaments composed of tubulin dimers. When one of the two flagella is sheared off at its base, the severed flagellum does not simply regrow to match the remaining one. Rather, as the severed flagellum begins to regrow, the longer flagellum shrinks until the two flagella are equal in size, after which both grow out in unison (1). If, as hypothesized, the limiting-pool mechanism is indeed the mechanism by which the flagellum lengths are made equal, then our results predict the existence of anti-correlated fluctuations in the lengths of the flagella as they swap tubulin dimers. Observations of these fluctuations would serve as a quantitative test of the limiting pool mechanism.

### Time scales of assembly

Earlier in this section, we discussed different time scales associated with growth of filaments from identical nucleating centers, i.e. growth and diffusion timescales and their dependence on the number of monomers in the pool. One can use those calculations to estimate timescales in the case where multiple actin cables are made in the same pool of actin monomers in the mother compartment of a budding yeast. Using previously published numbers for cell volume (11) and rates of association and dissociation of monomers to actin filaments (12), we estimate 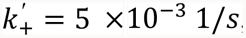, and number of actin cables (about ten), we predict that the growth phase lasts for less than a minute, assuming that there are no additional length control mechanisms. (See SI, section 2.1 and 3.5 for details).

In contrast, we estimate the diffusion time scale to span several days (SI, Section 2.3). In other words, we should never observe order-*N*^2^ fluctuations in cable lengths on the time scale of live-cell experiments, given a division time of about 1.5 hours. Note that for this estimate, we assume that all the actin in the mother compartment of the budding yeast cells are used to make cables; a reasonable assumption as these cells have very few or no patches in their mother compartment. A substantially smaller number of actin monomers in cables could bring down the estimate of the diffusion time scale considerably due to the *N*^2^ dependence of this time scale. The fact that large fluctuations in cable length are not observed could also be due the presence of additional length control mechanisms. In fact, several actin and formin binding proteins have been shown to play an important role in controlling cable length (5, 18, 19).

### Additional size-control mechanisms

In cells, multiple size-control mechanisms can work in unison to control the size of a single structure (3, 5, 4, 18, 19). The limiting-pool mechanism is very likely one of the mechanisms at play as cells usually contain a limited number of building blocks which are used to make internal structures. Since our results provide quantitative predictions regarding the sizes of structures assuming that the limiting pool mechanism is the only one at play, they can potentially be used to identify the existence of additional size-control mechanisms. For example, actin cables in budding yeast are estimated to be almost three times as long as those seen in experiments if the finite number of actin monomers were the only size controlling factor. And in fact a mechanism that utilizes directed transport of an assembly-attenuating factor by a myosin motor along the cable was shown to lead to a length dependent assembly rate, thereby limiting the size of these structures (5, 18).

In summary, the limiting-pool mechanism of size control can produce a single structure of a well-defined size. On the other hand, this mechanism is unable to maintain multiple structures that have a well-defined size, which assembly from a common pool of molecular components. Cells seem to get around this problem by using additional regulatory mechanisms, all of which seem to produce a length dependent assembly or disassembly rate. Quantitative experiments that measure the size of an intracellular structure and how they vary with changing amounts of diffusing components are needed to quantitatively define the role of the limiting-pool mechanism in regulating size.

## Material and methods

### Simulation protocol

We used stochastic simulations to solve the master equations in Equations 1, 2 and 4. We start with a filament of zero length and then follow the stochastic trajectory of the filament. In the simulation, the state of the system is characterized by the filament length. In one step of the simulation we choose one of the set of all possible transitions from the current state of the system to the next. The transitions are chosen at random according to their relative weight, which is proportional to the rate of the transition. Once a particular transition is chosen the system is updated to a new state, which becomes the new current state. The time elapsed between two consecutive transitions is drawn from an exponential distribution, the rate parameter of which equals the sum of all the rates of allowed transitions. This process is repeated for a long enough time such that the length of the cable reaches steady state. We obtain many such trajectories of a single filament and then compute the steady state distributions of filament lengths.

## Acknowledgements

We wish to thank Rob Phillips, Bruce Goode, Wallace Marshall, Stephanie Weber, David Kovar, Seham Ebrahim, Fred Chang and Nenad Pavin for many stimulating discussions. This work was supported by the National Science Foundation through grants DMR-1206146 and MRSEC-1420382 (J.K., L.M. and D.H.), Boehringer Ingelheim Funds, Arthur Klorfein Scholarship and Fellowship Fund, Mountain Memorial Fund Scholarship, Howard A. Schneiderman Endowed Scholarship (T.J.L). We are grateful to the Burroughs-Wellcome Fund for its support of the Physiology Course at the Marine Biological Laboratory, where part of the work on this paper was done.

## References

1. Goehring NW, Hyman AA (2012) Organelle growth control through limiting pools of cytoplasmic components. Curr Biol CB 22(9):R330–339.

2. Dumont S, Mitchison TJ (2009) Force and length in the mitotic spindle. Curr Biol CB 19(17):R749–761.

3. Avasthi P, Marshall WF (2012) Stages of ciliogenesis and regulation of ciliary length. Differ Res Biol Divers 83(2):S30–42.

4. Marshall WF, Qin H, Rodrigo Brenni M, Rosenbaum JL (2005) Flagellar length control system: testing a simple model based on intraflagellar transport and turnover. Mol Biol Cell 16(1):270–278.

5. Chesarone-Cataldo M, et al. (2011) The myosin passenger protein Smy1 controls actin cable structure and dynamics by acting as a formin damper. Dev Cell 21(2):217–230.

6. Michelot A, Drubin DG (2011) Building distinct actin filament networks in a common cytoplasm. Curr Biol CB 21(14):R560–569.

7. Good MC, Vahey MD, Skandarajah A, Fletcher DA, Heald R (2013) Cytoplasmic volume modulates spindle size during embryogenesis. Science 342(6160):856–860.

8. Weber SC, Brangwynne CP (2015) Inverse Size Scaling of the Nucleolus by a Concentration-Dependent Phase Transition. Curr Biol 25(5):641–646.

9. Buttery SM, Yoshida S, Pellman D (2007) Yeast formins Bni1 and Bnr1 utilize different modes of cortical interaction during the assembly of actin cables. Mol Biol Cell 18(5):1826–1838.

10. Johnston AB, Collins A, Goode BL (2015) High-speed depolymerization at actin filament ends jointly catalysed by Twinfilin and Srv2/CAP. Nat Cell Biol 17(11):1504–1511.

11. Philips RM& R Cell Biology by the Numbers. Available at: http://book.bionumbers.org/ [Accessed June 24, 2016].

12. Pollard TD (1986) Rate constants for the reactions of ATP- and ADP-actin with the ends of actin filaments. J Cell Biol 103(6 Pt 2):2747–2754.

13. Kelly, F. P. Reversibility and Stochastic Networks; Cambridge University Press, 2011.

14. Hu J, Othmer HG (2011) A theoretical analysis of filament length fluctuations in actin and other polymers. J Math Biol 63(6):1001–1049.

15. Brugués J, Nuzzo V, Mazur E, Needleman DJ (2012) Nucleation and Transport Organize Microtubules in Metaphase Spindles. Cell 149(3):554–564.

16. Suarez C, et al. (2015) Profilin regulates F-actin network homeostasis by favoring formin over Arp2/3 complex. Dev Cell 32(1):43–53.

17. Rotty JD, et al. (2015) Profilin-1 serves as a gatekeeper for actin assembly by Arp2/3-dependent and -independent pathways. Dev Cell 32(1):54–67.

18. Mohapatra L, Goode BL, Kondev J (2015) Antenna Mechanism of Length Control of Actin Cables. PLoS Comput Biol 11(6):e1004160.

19. Mohapatra L, Goode BL, Jelenkovic P, Phillips R, Kondev J (2016) Design Principles of Length Control of Cytoskeletal Structures. Annu Rev Biophys. doi:10.1146/annurev-biophys-070915-094206.

20 Ishikawa H, Marshall WF (2011) Ciliogenesis: building the cell’s antenna. Nat Rev Mol Cell Biol 12(4):222–234.

